# The Evolutionary Trade-off between Stem Cell Niche Size, Aging, and Tumorigenesis

**DOI:** 10.1101/059279

**Authors:** Vincent L. Cannataro, Scott A. McKinley, Colette M. St. Mary

## Abstract

Many epithelial tissues within large multicellular organisms are continually replenished by small independent populations of stem cells. These stem cells divide within their niches and differentiate into the constituent cell types of the tissue, and are largely responsible for maintaining tissue homeostasis. Mutations can accumulate in stem cell niches and change the rate of stem cell division and differentiation, contributing to both aging and tumorigenesis. Here, we create a mathematical model of the intestinal stem cell niche, crypt system, and epithelium. We calculate the expected effect of fixed mutations in stem cell niches and their expected effect on tissue homeostasis throughout the intestinal epithelium over the lifetime of an organism. We find that, due to the small population size of stem cell niches, fixed mutations are expected to accumulate via genetic drift and decrease stem cell fitness, leading to niche and tissue attrition, and contributing to organismal aging. We also explore mutation accumulation at various stem cell niche sizes, and demonstrate that an evolutionary trade-off exists between niche size, tissue aging, and the risk of tumorigenesis; where niches exist at a size that minimizes the probability of tumorigenesis, at the expense of accumulating deleterious mutations due to genetic drift. Finally, we show that the probability of tumorigenesis and the extent of aging trade-off differently depending on whether mutational effects confer a selective advantage, or not, in the stem cell niche.

## 1 Introduction

The constant occurrence of errors in DNA replication, coupled with the imperfect nature of damage recognition and repair mechanisms, results in the continual accumulation of mutations in the genome within the cells constituting multicellular organisms [Lynch, 2008]. The full extent of mutation accumulation, and the burden these mutations place on the somatic tissues within multicellular organisms, is not known. Here, we utilize the recently elucidated population dynamics of intestinal stem cells, along with the dynamics of stem cell progeny, to model the evolutionary tendency of stem cell populations and the expected burden that mutation accumulation places throughout the intestinal tissue. The paradigm of stem cell turnover within the intestine is not unique to that tissue [Klein and Simons, 2011], and thus the work presented here may be applicable to other systems undergoing continual self-renewal through the division of stem cells.

**Fitness and Aging**. The majority of mutations that affect an individual’s fitness will decrease fitness [Eyre-Walker and Keightley, 2007]. In populations of whole organisms, especially those in which fitness effects are studied in the laboratory, these mutations rarely become fixed due to purifying selection and large population sizes (see Levy et al. [2015] for a recent example). However, many of the independently evolving populations of adult stem cells that continually divide to replenish tissues are maintained at very small population sizes [Morrison and Spradling, 2008], and are thus more susceptible to genetic drift and the fixation of deleterious mutations. In addition, due to the asexual nature of mitotic division and stem cell maintenance, stem cell populations are a prime example of Muller’s ratchet, or, the irreversible accumulation of mutations and resultant decrease in a population’s mean fitness in the absence of recombination [Muller, 1964, Lynch et al., 1993]. We have previously demonstrated that crypts are predominantly accumulating deleterious mutations over the lifetime of the host organism [Cannataro et al., 2016]. Here, we quantify the expected change in whole-tissue equilibrium population size as mutations accumulate in the stem cell populations at the base of intestinal crypts over an organism’s lifetime.

The accumulation of damage causing the loss of cellular fitness is a hallmark of aging, and is especially relevant when DNA damage occurs in stem cells, compromising their role in tissue renewal [Lopez-Otln et al., 2013]. Indeed, several mouse models with the diminished ability to maintain cellular genome integrity succumb to accelerated age-related phenotypes through the loss of tissue homeostasis caused by stem and progenitor cell attrition [Ruzankina et al., 2008]. Just as stem cell mutations conferring a beneficial fitness effect will increase cell production, mutations conferring a deleterious fitness effect will lead to decreased cell production and the diminished maintenance of healthy tissue. Stem cells at the base of the intestinal crypt differentiate into all other intestinal cell populations [Barker, 2014]. Hence, mutations affecting the rates of stem cell dynamics will propagate through other populations, affecting their steady state equilibrium population sizes. We model the various cell populations of the intestinal crypt and epithelium to calculate how mutations occurring in stem cell lineages govern population dynamics throughout the tissue.

## 2 Methods/Modeling

We are interested in how the rates of stem cell division and differentiation scale up to whole tissue dynamics. Within this section we first describe the general architecture of crypt systems. Then, we detail our parameterization of the models in light of recent experiments in mice. Finally, we describe how we model the evolutionary processes occurring within crypts and throughout the tissue.

### 2.1 Modeling the Crypt System

**Intestinal crypts are composed of the various types of cell populations derived from stem cells**. Mouse crypts contain a population of LGR5^+^ putative stem cells at their base. Within this population, there exists a functional subpopulation responsible for maintaining homeostasis within the crypt [Kozar et al., 2013, Baker et al., 2014]. We call this subpopulation responsible for crypt homeostasis *X*_1_. The stem cells in the stem cell niche divide symmetrically and undergo neutral drift, where any lineage with the same division rate as the other lineages in the crypt has equal probability to reach monoclonality by displacing all other lineages through division [Ritsma et al., 2014]. Cells with greater division rate have an increased probability of displacing their neighbors; those with lower division rate are more likely to be displaced. In our model, both stem cells within the niche and displaced stem cells divide symmetrically at rate λ, and displaced stem cells commit to differentiation at rate *ν* and join the Transient-Amplifying (TA) compartment, *Y*_1_, located above the stem cell compartment. A TA cell rapidly divides at rate *γ* a number of rounds, *R*, joining subsequent TA pools (*Y*_1_,*Y*_2_, …,*Y_R_*). After the last round the cell divides a final time and joins the terminally-differentiated postmitotic cell pool *Z* [Potten, 1998]. Cells within the post-mitotic cell pool exist until they undergo apoptosis at rate *δ* either at the villus tip or lumenal surface in the small intestine and large intestine, respectively [Grossmann et al., 2002]. The terminally differentiated cells maintain the functionality of the intestinal tissue, with many existing at the top of the crypt, on the epithelial surface lining the lumen, and, in the case of the small intestine, along the villi. The dynamics described above are depicted in Figure 1.

These dynamics are represented by the transition rates

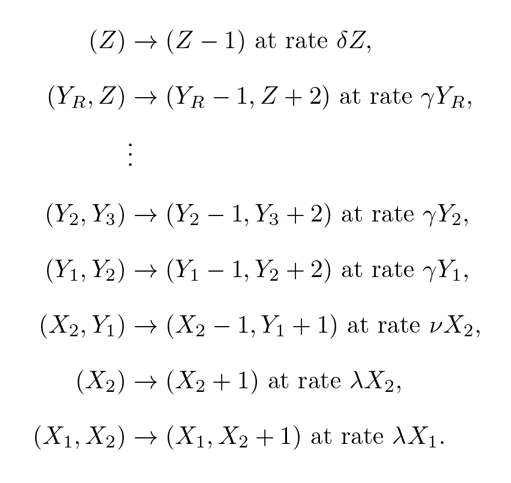

The time-dependent means of this system satisfy the ordinary differential equations:

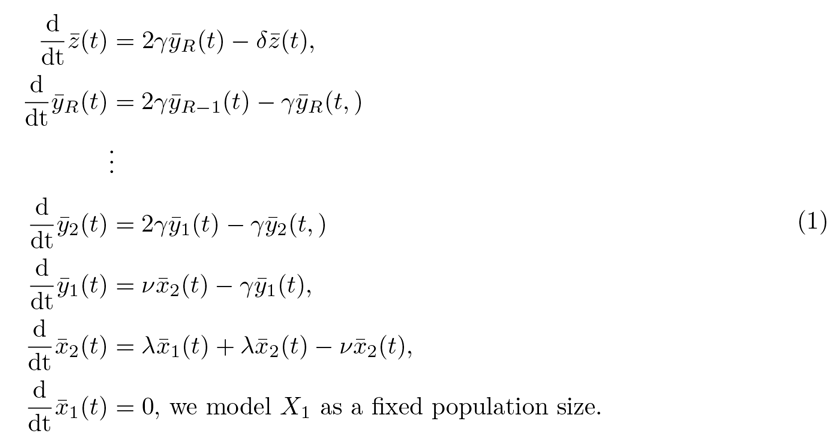

**Figure 1:**
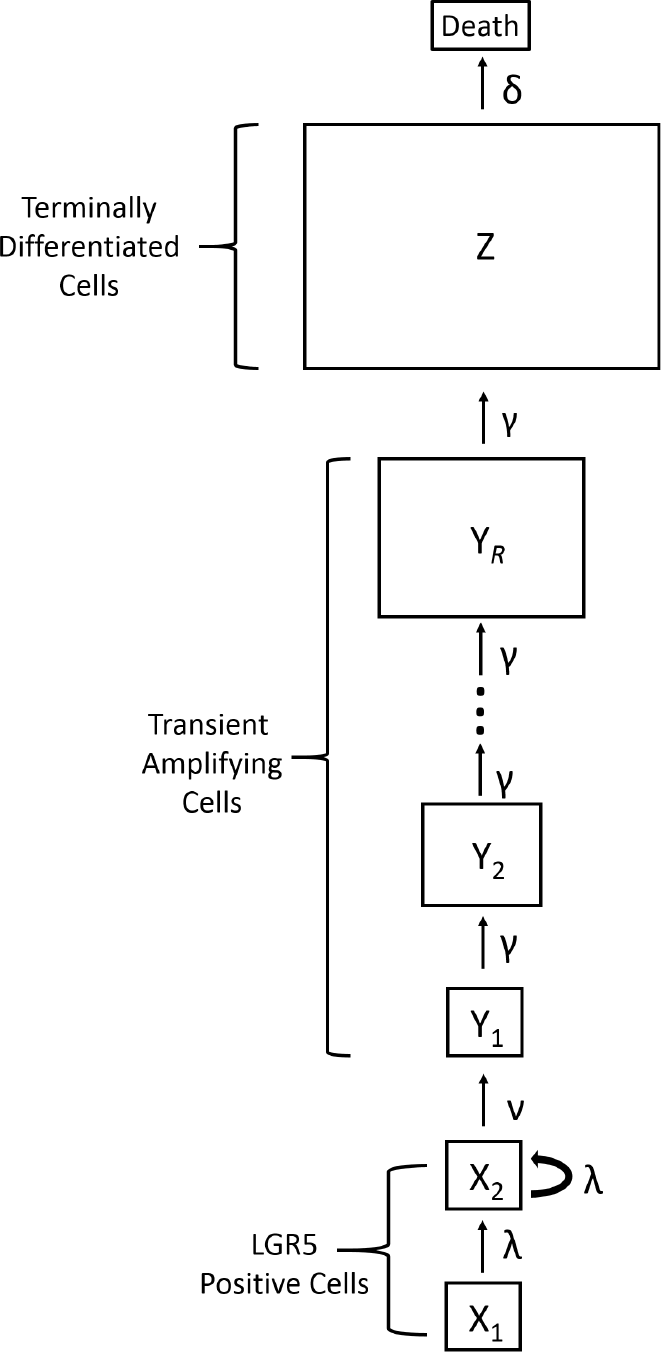
The general architecture of a crypt system. Population names are within the boxes and the rates at which cells accumulate within or are transferred between populations are next to the arrow portraying their transition.

Setting the left-hand sides for each equation in the system (1) to zero, we can solve for the steady state mean of the terminally differentiated population, *Z*\*. We find that *Z*\* can be expressed in terms of the system’s rate parameters and the stem cell niche population size *X*_1_. The steady state *Z*\* (Eq. 2) is a function of all of the rate parameters in the system except the TA cell division rate:

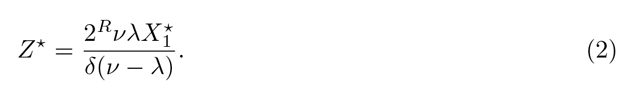

This function is used in subsequent analyses to calculate how mutations to the various rate parameters change the tissue population size. Of note, a prediction of this model is that a mutation to either division rate or differentiation rate will result in an amplified effect on the proportion of change in steady state post-mitotic cells. That is, if λ_0_ is mutated to λ_1_, the proportional difference in the post-mitotic cell population is 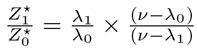.

**Table 1:** Approximate parameter values for the mouse intestine

| Parameter | Value | Note | Citation |
| --- | --- | --- | --- |
| $X_1$ | 6 Cells | Expected number of stem cells in the small intestine stem cell niche | Kozar et al. [2013] |
| $X_1 + X_2^*$ | 15 Cells | The total number of Lgr5<sup>+</sup> in the base of the crypt | Snippert et al. [2010], Clevers [2013] |
| $\lambda$ | 0.2 per day | The division rate of the sub population of the stem cells constituting the niche | Kozar et al. [2013] |
| $\nu$ | 0.333 per day | The rate at which cells passively adopt a TA fate | this study |
| $\gamma$ | 2 per day | The division rate of TA cells | Potten and Loeffler [1987] |
| $\delta$ | 0.333 per day | The death rate of terminally differentiated cells | Snippert et al. [2010] |
| $R$ | 6 | Rounds of division of rapidly dividing TA cells | Marshman et al. [2002] |

### 2.2 Model Parameterization

**Parameterizing the crypt model using mouse small intestine data**. The most studied crypt system is in the mouse small intestine. There are 14-16 Lgr5^+^ putative stem cells in the base of the mouse small intestinal crypts and it is believed that they divide symmetrically under neutral competition [Snippert et al., 2010, Lopez-Garcia et al., 2010]. Kozar et al. [2013] found that there exists a subset of this population that maintains crypt homeostasis, and that this subpopulation consists of approximately 5, 6, and 7 cells dividing approximately 0.1, 0.2, and 0.3 times per day along the proximal small intestine, distal small intestine, and colon, respectively. When estimating the expected effects of mutation accumulation along the entire intestine in the mouse we take the median value of cell number and division rate found by Kozar et al. [2013] to be the number of cells in the niche, *X*_1_, and the stem cell symmetric division rate, λ, with the remainder of the stem cell population constituting the *X*_2_ population. In order to maintain the number of Lgr5^+^ stem cells at a steady state of 15 we calculate a rate of committing to differentiation *ν* of 0.333 per day. Transient amplifying cells divide twice a day [Potten and Loeffler, 1987, Potten, 1998] and undergo approximately six generations of transient amplification, with each stem cell eventually contributing about 64 cells to the crypt system [Marshman et al., 2002]. These lineages end at their terminally differentiated cell stage, and live for 2-3 days before dying and leaving the intestine [Snippert et al., 2010]. The parameters described above are provided in Table 1.

Parameterizing our structured model with the values from Table 1 the model generates observed cell pool population sizes. Using the model we calculate the steady state mean population sizes of the various TA populations to be 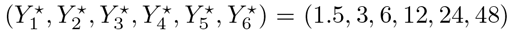. This results in the total rapidly dividing TA population being 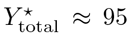 cells, slightly underestimating estimates from the literature of the number of cells within this compartment which are around 120 [Marshman et al., 2002]. These dynamics result in a steady state mean of the terminally differentiated cell population size in our model, *Z*\*, to equal 576 cells. The crypts consist of approximately 250 cells [Potten and Loeffler, 1990], meaning that (accounting for the 95 TA cells, the 15 stem cells, and the 10 Paneth cells not modeled here [Clevers, 2013]) the terminally differentiated cell population in our model exists as 130 cells within the crypt and 466 cells outside of the crypt. The villi in the small intestine are supported by 6-10 crypts and consist of approximately 3500 cells [Potten and Loeffler, 1990]. The total steady state mean population in our model contributed by 8 crypts to a villus using the parameters in Table 1 is 8 × 446 = 3568 cells.

We are interested in estimating the effects of mutation accumulation throughout the whole intestinal tract in the mouse, hence we take the median value of certain parameters that vary from proximal small intestine to colon (as described above). We note that if one was interested in the effects of steady state terminally differentiated population size in just the proximal small intestine or colon this analysis may overestimate or underestimate the effects, respectively, since there are more stem cells dividing faster in the colon than the proximal small intestine. Additionally, the large intestine TA cells may undergo more rounds of division than the small intestine TA cells [Potten, 1998].

**Intrinsic mutational effects**. The entire distribution of fitness effects of mutations is unknown for somatic tissues. Additionally, although the mutation rate has been estimated on a per nucleotide basis [Jones et al., 2008], the mutational target size of all mutations that affect crypt cell fitness is unknown. Thus, we explore the implications of a wide range of intrinsic mutational parameters within somatic tissue. However, despite the differences between the genomes of species, mutation accumulation experiments and analysis of DNA sequence data have revealed general principles regarding distributions of intrinsic mutational effects. Namely, mutations are much more likely to confer a deleterious fitness effect than a beneficial effect, and that the deleterious effect will have a larger expected value [Eyre-Walker and Keightley, 2007, Bank et al., 2014]. Deleterious fitness effects are extremely difficult to characterize in typical laboratory experiments because selection against deleterious mutations is effective in even moderately sized populations [Estes et al., 2004, Keightley and Halligan, 2009]. Directed mutagenesis experiments are one way to classify the distribution and average effect of mutations deleterious to fitness. For instance, Sanjuan et al. [2004] performed site-specific single-nucleotide substitutions in an RNA virus and found that the average deleterious non-lethal fitness effect decreased fitness approximately 20%. Similarly, when grown in permissive conditions that allow spontaneous deleterious mutations to accumulate through genetic drift, Keightley and Caballero [1997] found a 21% average deleterious fitness effect per mutation in *Caenorhabditis elegans* and Zeyl and DeVisser [2001] found a 21.7% average fitness decline per fixed mutation in diploid strains of the single celled eukaryote *Saccharomyces cerevisiae.* Other mutation accumulation studies have found more modest deleterious effects, such as the average fitness decline in haploid *Saccharomyces cerevisiae* per mutation of 8.6% found by Wloch et al. [2001]. Another mutation accumulation experiment in *Saccharomyces cerevisiae* found the expected beneficial increase of fitness per mutation to be 6.1%, the rate of mutation that affects fitness per mutation to be 1.26 ×10^−4^, and the percent of fitness effects that are beneficial to be 5.75% [Joseph and Hall, 2004]. When our analysis requires specific parameter choices, as in Section 3.3 when we juxtapose the dynamics of mutations that fix neutrally with those under selection, we utilize the *Saccharomyces cerevisiae* parameters described here, but note that we are interested in characterizing the dynamics of tumorigenesis and aging, and we are not making conclusions about the absolute magnitude of either given the limited knowledge of mutational effects in somatic tissue.

### 2.3 Modeling Evolution Within Somatic Tissue

**Modeling the expected mutational effect of a single mutation within a crypt**. In order to quantify the expected effect on tissue homeostasis of mutations in epithelial tissue it is necessary to understand the processes of mutation accumulation and fixation within the stem cell niche populations at the base of the intestinal crypts. Mutations in the niche can be placed into two different categories: mutations that directly affect the stem cell phenotype associated with cellular fitness, i.e. division rate, within the stem cell niche, and mutations that do not affect the fitness of stem cells within the niche. Mutations that affect the division rate of stem cells will confer a fitness advantage or disadvantage because stem cells within the niche are symmetrically dividing and replacing their neighbors. For instance, certain mutations to KRAS increase stem cell division rate and the probability this mutant lineage reaches fixation [Vermeulen et al., 2013, Snippert et al., 2014]. Mutations that do not directly affect stem cell division rate will not alter stem cell fitness within the niche and will fix neutrally.

We model the distribution of mutational effects and mutation accumulation similarly as in Cannataro et al. [2016], where we provide a detailed mathematical methodology. Briefly, mutational effects are distributed exponentially, with expected deleterious effect *s*_, expected beneficial effect *s*_+_, and *P_B_* percent of mutations conferring a beneficial effect. Thus, mutational effects on the initial stem cell division rate, λ_0_, are distributed according to Eq. 3, where

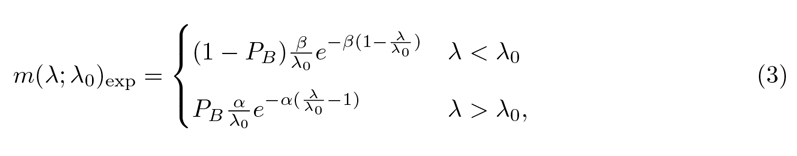

where 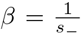 and 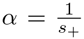. Distributions that are more leptokurtic than exponential, and/or bimodal, may provide better characterization of deleterious and beneficial mutations [Eyre-Walker and Keightley, 2007, Levy et al., 2015], however, modeling the DFE as an exponential distribution minimizes the number of parameters in our assumptions while still capturing a distribution that has provided reasonably good fits to both deleterious [Elena et al., 1998] and beneficial [Kassen and Bataillon, 2006] fitness effects during experiments.

A new lineage with a division rate relative to the background division rate 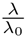 has probability of eventually replacing the original lineage

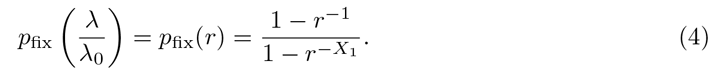

We use Bayes’ theorem to calculate the probability density of the division rate given the mutant lineage fixed,

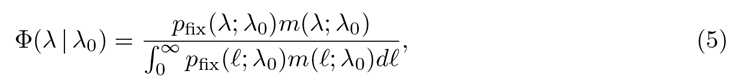

and, redefining Eq. 5 such that *f*_1_(λ) is equal to the density given the first fixation of a mutant lineage, we calculate the expected value of the division rate of this new lineage: 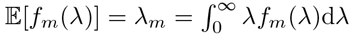.

Note that we can extend this analysis to the accumulation of mutations that do not alter the fitness of stem cells within the niche by changing Eq. 4 to be equal to 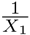, the probability of fixation of a neutral mutation. Additionally, mutations that affect differentiation rate can also be modeled using Eq. 3 by switching the direction of effect such that beneficial mutations now decrease the differentiation rate (i.e. increase the lifetime of the cells).

**Calculating the expected effect of multiple mutations within a crypt, and the accumulation of mutations throughout all crypts**. We calculate probability densities describing the division rate of *m* subsequent fixed mutations, and the probability of tumorigenesis they confer, according to the recursive formula

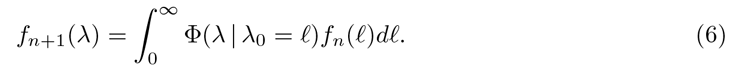

From Eq. 6 we can calculate the expected value of division rate given m mutations, which is the expected value of these probability densities: 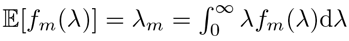.

We model the rate at which new lineages arise and fix in the crypts as being constant over time with rate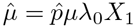 where 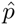 is the total probability of fixation 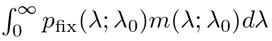 and *μ* is the mutation rate, as in Cannataro et al. [2016]. Here, the number of fixed mutations within a crypt, *m*, are approximated by the Poisson distribution with mean 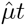, and we can estimate the number of *n* crypts with *m* mutations by multiplying this distribution by the number of crypts in the system, *C*:

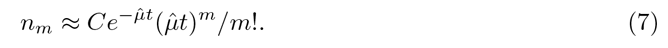

Thus, we can estimate the total number of crypts with m mutations as the organism ages.

**Calculating the probability that fixed mutations initiate tumorigenesis**. For part of our analysis we juxtapose the expected magnitude of tissue change with the risk of tumorigenesis for a given stem cell niche size. We define the initiation of tumorigenesis as the moment when the division rate of stem cells, λ, becomes greater than the differentiation rate of stem cells, *ν*, thus initiating exponential growth within the tissue. We calculate this by integrating the probability densities describing the change in the stem cell rates from *ν* to infinity for scenarios where mutations affect division rate, and λ to zero for scenarios where mutations affect differentiation rate, with both calculations determining the probability that a certain mutation resulted in a fitness change that initiated tumorigenesis. These probabilities are summed over all mutations in all crypts, giving the total probability of tumorigenesis.

**Calculating the expected effect of mutation accumulation on tissue maintenance**. Using our calculation of the number of crypts with *m* mutations (Eq. 7), the expected division rate of stem cells within niches with *m* mutations, and the steady state post-mitotic cell population given this expected division rate (Eq. 2), we can estimate the expected post-mitotic epithelial population size over the time since adulthood, *t*, of the individual,

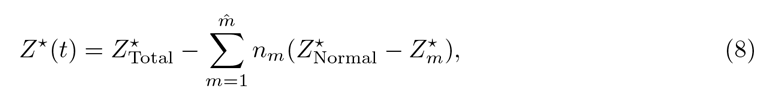

where 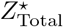 is equal to the total number of crypts times the number of post-mitotic cells produced by a crypt with no fixed mutations, 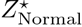.

Mutations that do not affect stem cell division rate fix neutrally in the stem cell niche, however, these mutations may still alter the steady state mean of the post-mitotic terminally differentiated population by changing population dynamics rates that are important for tissue maintenance. For instance mutations that strictly deal with the rate for a lineage to differentiate (*ν*) would still change *Z*\*.

### 2.4 Linear Approximation

Plots of 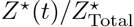 as a function of organismal lifetime are approximately linear (see Section 3.2). Hence, we utilize an asymptotic analysis to approximate the changes to the post-mitotic cell population over a lifetime, i.e., the dynamics defined by Eq. 8. When only considering the relative affect of the first fixed mutation within stem cell niches, and the arrival of this first fixed mutation among the niches, we can simplify Eq.8 to

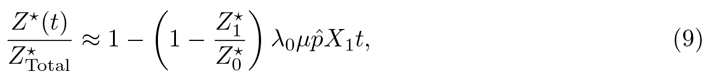

where 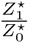 is the steady state population size of the post-mitotic cell population after one fixed mutation divided by the healthy population size with zero mutations (Eq. 2). When mutations alter the division rate the fraction 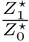 simplifies to 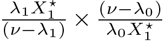 or 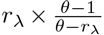, where 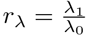 and 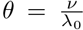. Thus, Eq. 9 simplifies to 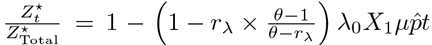 whose time derivative is

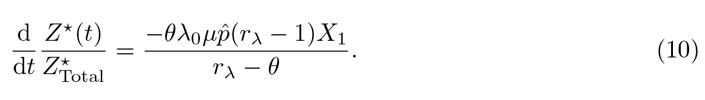

When mutations affect the differentiation rate of stem cells, the fraction 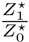 simplifies to *r*_*ν*_ × 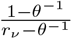, where 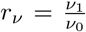. In this case Eq. 9 simplifies to 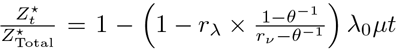, and the derivative of this function with respect to time is:

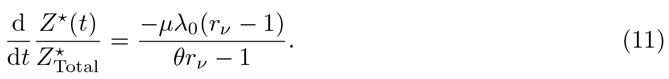

Note how Eq. 11 is independent of *X*_1_ and 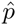, since mutations fix neutrally and these terms are the inverse of one another.

Equations 10 and 11 are close approximations of the rate of tissue size change per day. Importantly, they are independent of many of the parameters in the crypt model (1), such as the division rate and number of generations of TA cells and the apoptosis rate of post-mitotic cells. These parameters are not easily measured in humans, and thus Equations 10 and 11 enable us to extend our inference to human scenarios.

**Parameterization for the human colon**. The division rate of stem cells within human colon crypts has been has been reported as close to once a week [Potten et al., 2003, Nicolas et al., 2007] and the total number of stem cells maintaining the crypt population, i.e. stem cells in the niche, has been estimated to be between 6 [Baker et al., 2014] and 15 to 20 cells [Nicolas et al., 2007], with Bayesian analysis from the latter study showing more support for larger crypt population sizes. Assuming the human stem cell niche in the colon crypt houses 6 cells, and the total mean number of potential stem cells in the colon crypt is approximately 36 cells [Bravo and Axelrod, 2013], we use Equation 1 to calculate that the differentiation rate of stem cells is approximately 0.172 per day. Alternatively, if the human stem cell niche houses 20 cells, and there are a total of 36 putative stem cells within the crypt, we calculate the differentiation rate of stem cells to be 0.321 per day. Thus, we have all the parameters necessary to calculate the expected effect of the accumulation of mutations in stem cell niches on tissue homeostasis in human intestines, and can compare the expected effects given the two different estimated sizes of the stem cell niche.

### 2.5 Exploring the evolutionary trade-off between stem cell niche size, aging, and tumorigenesis

Given that small asexual populations are prone to succumb to a gradual decline in mean fitness via the accumulation of deleterious mutations [Lynch et al., 1993] we explore the selective pressures that may have influenced the evolution of small stem cell niche population size. Specifically, we juxtapose the magnitude of epithelium tissue population change and the total risk of tumorigenesis throughout the intestinal epithelium for different stem cell niche sizes. Throughout this analysis we assume that half of the putative stem cells in the crypt reside in the niche, as is approximately the case for both mice and humans (see above). Additionally, we emphasize that the absolute magnitudes of tissue change and tumor incidence may not reflect the true values, as the parameters associated with the true expected mutational effects are unknown,however the emergent selective dynamics from our analysis are relatively independent of exact parameter specifications.

## 3 Results

### 3.1 The expected fitness of stem cells decreases with fixed mutations

**When mutations affect division rate**. The small population size of the stem cell niche at the base of the crypts promotes weak selection and pervasive drift throughout the intestine. For each crypt, across a range of distribution of fitness effect parameters consistent with those measured in whole organisms and stem cell niche sizes consistent with those measured in mice and humans (between 6 and 20 cells), the expected value of the first fixed mutation has a lower fitness than the previous lineage when mutations affect division rate (Figure 2). The expected value of a fixed mutation that affects division rate increases with larger values of population size, the expected effect of a beneficial mutation, and the probability that a mutation results in a beneficial effect.

**Figure 2:**
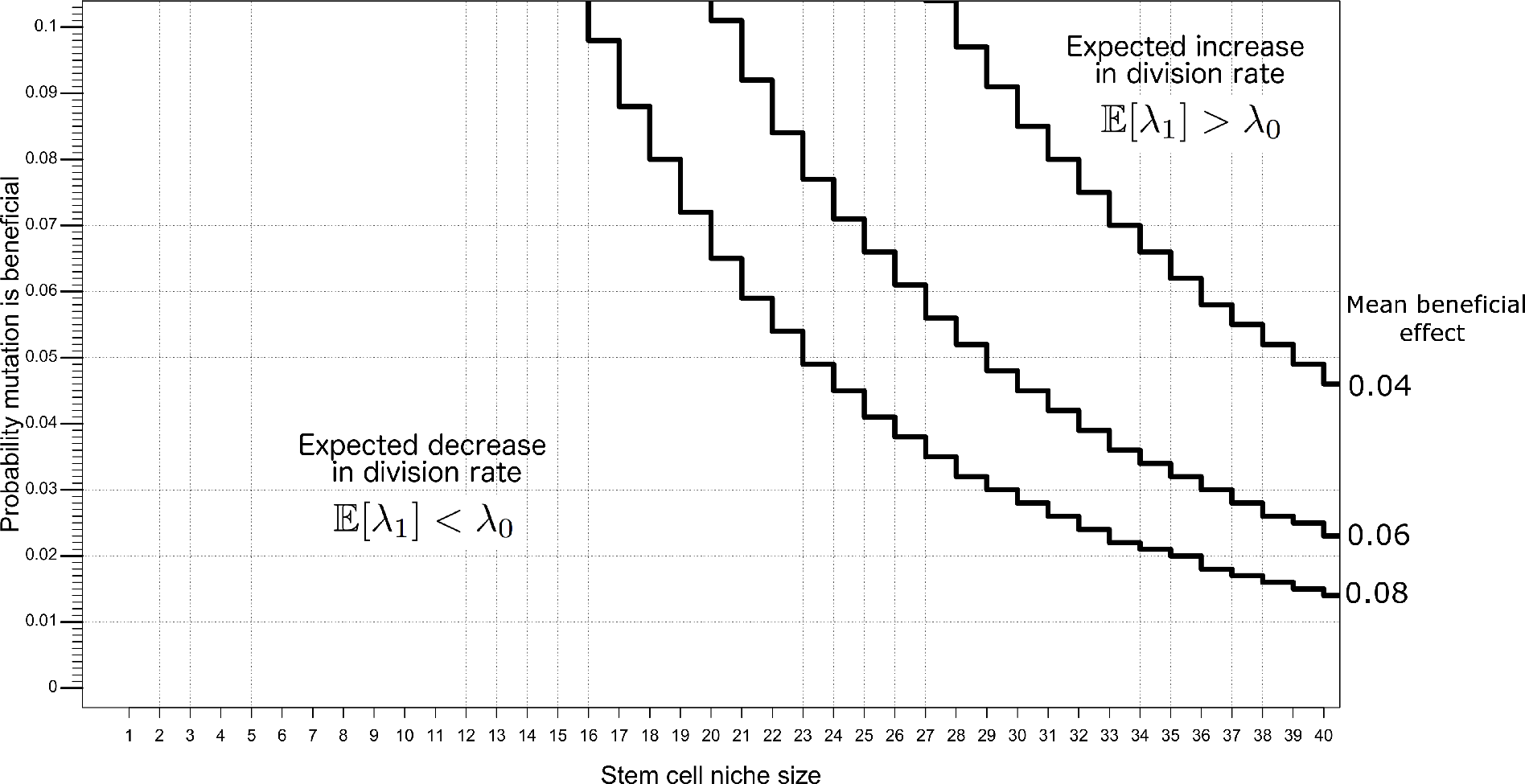
Expected values of division rate after a fixed mutation are lower than the original division rate (deleterious) for reported stem cell niche sizes and plausible DFE parameters. The lines plotted here represent the boundary in the parameter space that separates the scenarios where beneficial or deleterious mutations are expected to accumulate. The mean deleterious effect for the parameter space depicted here is 0.15.

The power of genetic drift at small populations is exhibited by considering the expected effect of a fixed mutation as a function of the expected effect of deleterious fitness effects at two population sizes (Figure 3). For smaller populations, such as when *X*_1_ = 6, even mutations of large deleterious effect may fix through genetic drift, and then the expected value of the first fixed mutation continues to decrease as the expected value of a deleterious mutation increases. Alternatively, larger stem cell niche population sizes are less prone to deleterious mutations fixing through drift, and as the expected effect of deleterious mutations increase, there is a point at which deleterious mutations become less and less likely to fix. Given a fixation event, it is likely the mutation conferred a beneficial effect.

**Figure 3:**
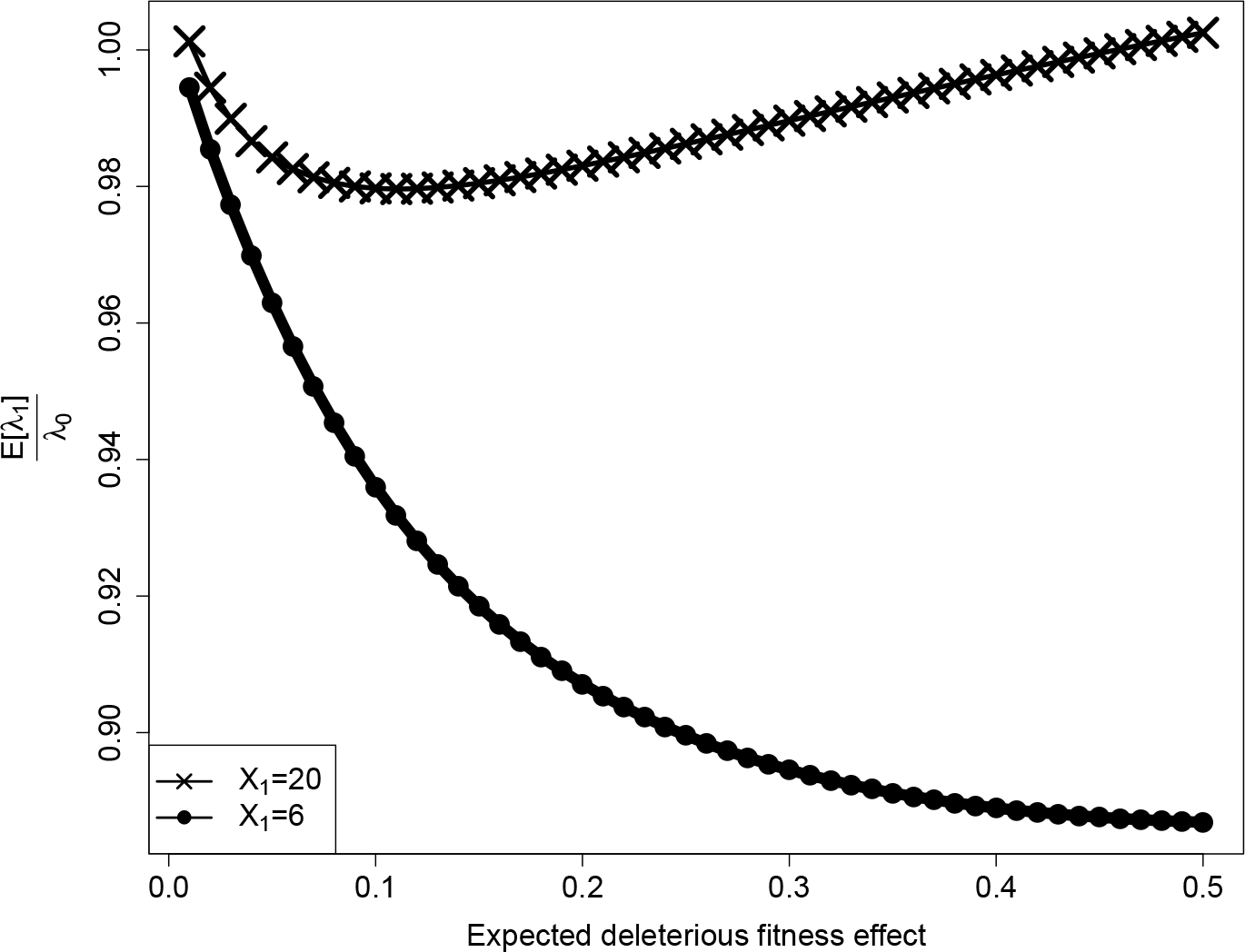
The expected value of division rate divided by the original division rate, *r*_λ_, versus *s_-_*, the expected mutational effect given a deleterious mutation, for the stem cell niche sizes of 20 and 6. Here, s_+_ is 0.061 and *P_B_* is 0.0575.

**When mutations affect differentiation rate**. Since the differentiation rate phenotype is not expressed in the niche, all mutations to this rate fix neutrally, and the expected value of *ν* given a fixed mutation does not depend on niche population size. Because the vast majority of mutations are deleterious, the expected value of mutations is always deleterious in the absence of selection.

### 3.2 Mouse and Human Stem Cell Fitness is Expected to Decrease with Age, Reducing Tissue Renewal

As demonstrated in section 3.1, the fitness of stem cells is expected to decrease as mutations accumulate in stem cell niches. This results in diminished whole-tissue population sizes as mutations accumulate throughout the crypts within the tissue in mice (Figure 4) and humans (Figure 5). We find that the linear approximation described in section 2.4 provides a good approximation to the simulated curves and we thus employ this approximation in estimating tissue size change curves for humans. Interestingly, when considering mutations that affect division rates in mice and humans (assuming a human stem cell niche population size of 20), we predict similar post-mitotic cell population size changes throughout organism lifetimes, with population size declining approximately 0.35% and 0.5% respectively. If we assume that the population size in the human stem cell niche is 6 and mutations only affect division rate we predict a decline in population size of approximately 12% over human lifetime. When mutations only affect differentiation rate we predict larger declines in population size, as there is no selective pressure against the fixation of deleterious mutations in this scenario.

**Figure 4:**
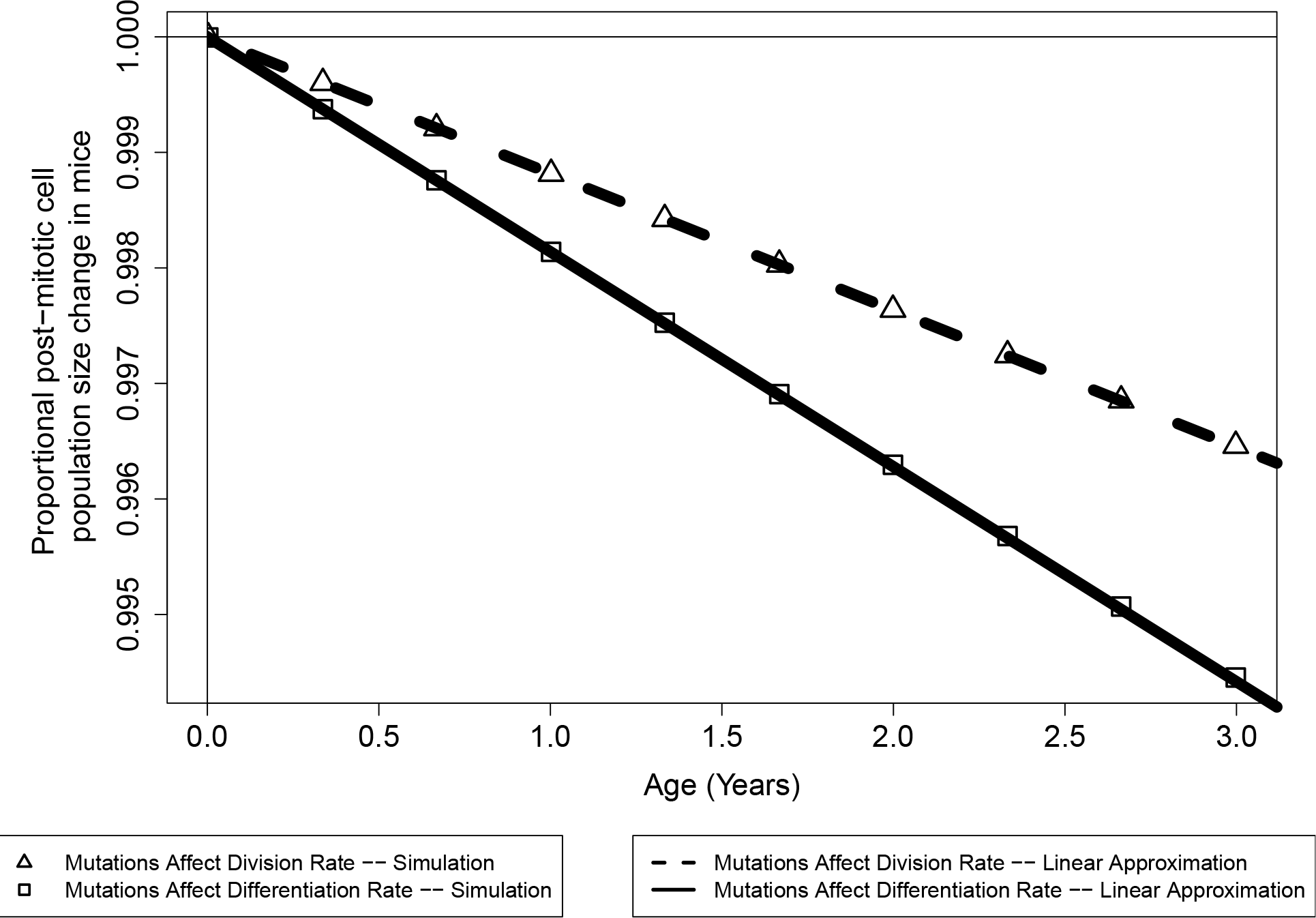
The expected tissue size change in mice due to mutation accumulation.

**Figure 5:**
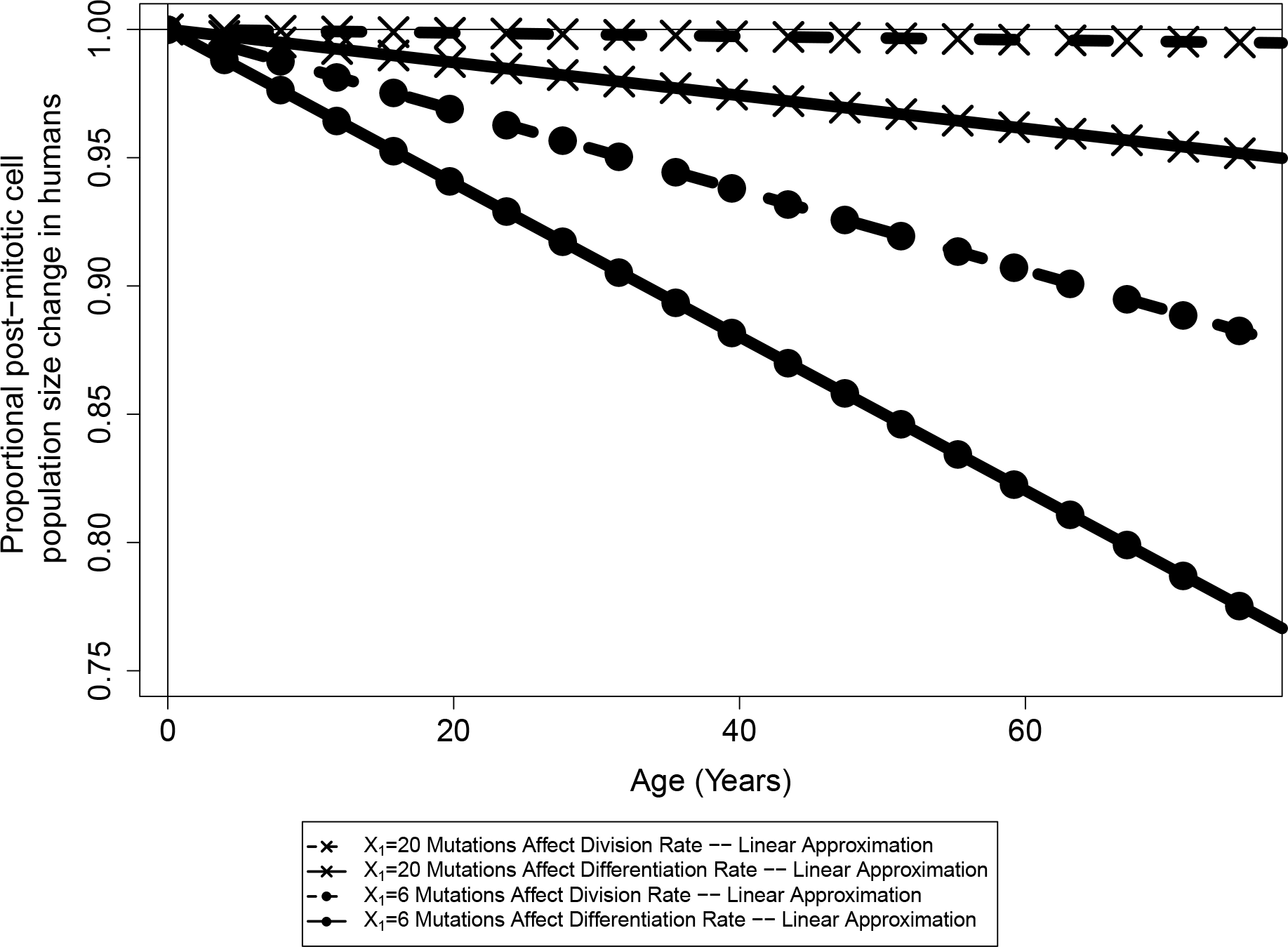
The expected tissue size change in humans due to mutation accumulation.

### 3.3 There is an evolutionary trade-off between tumorigenesis and aging mediated by stem cell niche size

We vary initial stem cell population size, along with the total number of crypts in the system, such that the total output of crypts, i.e., healthy tissue, remains constant. This analysis was initially conducted using the crypt parameters described for the mouse in Table 1 and the yeast mutation parameters described in section 2.2, namely mutations that affect fitness occur at rate 1.26 × 10^−4^, beneficial fitness effects have an expected value of 0.061, deleterious effects have an expected value of 0.217, and 5.75 percent of mutations confer a beneficial fitness effect. For mutations that affect division rate, there exists an optimal intermediate crypt size to minimize the probability of tumorigenesis (Figure 6A). However, at this crypt size, the expected value of the epithelium tissue size is expected to decrease over a lifetime due to the accumulation of deleterious mutations in stem cell niches (Figure 6B). Furthermore, when mutations affect differentiation rate and fix neutrally, the probability of tumorigenesis is minimized for large stem cell niche sizes (Figure 6C) and the expected effect on tissue size is invariant to stem cell niche size (Figure 6D). As the true mutational parameters governing somatic tissue evolution are unknown, we do not wish to analyze the predicted magnitude of tumorigenesis and aging presented here, but rather the trade-off that exists between niche population size, drift, and selection among mutations that confer a selective advantage and those that fix neutrally. The nature of the dynamics presented in Figure 6 holds true for a large range of mutation parameters. For instance, we next analyzed this evolutionary trade-off when considering the parameters discussed in Section 2.4 for the human colon (Figure 7). We find that the probability of a tumorigenesis event is minimized at a similarly small population size when mutations affect division rate, and there is a higher probability of tumorigenesis and larger change in population size for all human scenarios when compared to the results from the mouse model.

**Figure 6:**
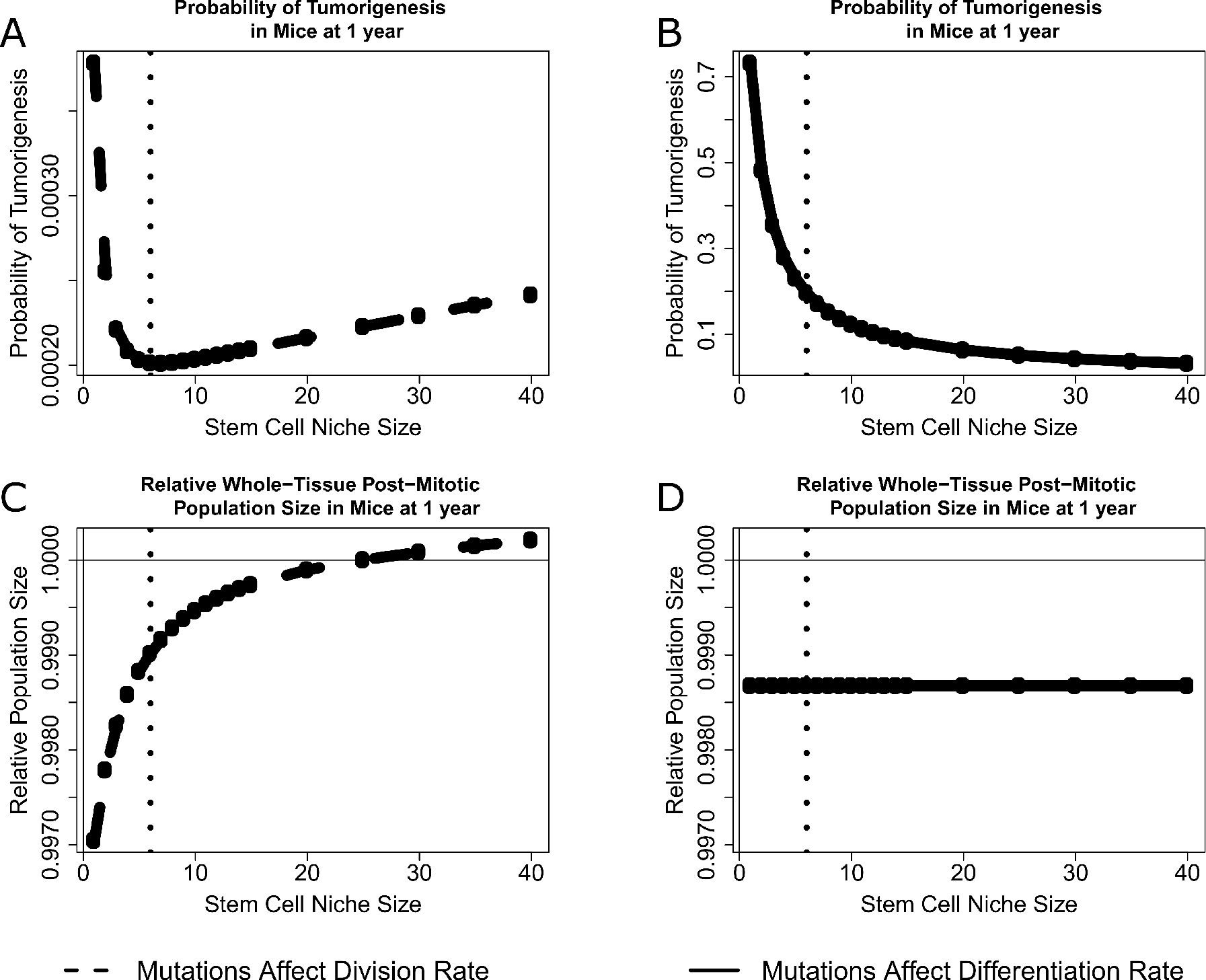
An Evolutionary Tradeoff with Niche Size in Mice. For scenarios where mutations affect division rate, tumorigenesis is minimized at an intermediate population size (A), at the expense of fixing deleterious mutations and decreasing tissue renewal (C). For scenarios where mutations affect differentiation rate, tumorigenesis is minimized for large population sizes (B), and the effect of mutation accumulation on the epithelium is independent of niche size (D). The black vertical dotted line is placed at 6, the median niche size measured in the mouse.

**Figure 7:**
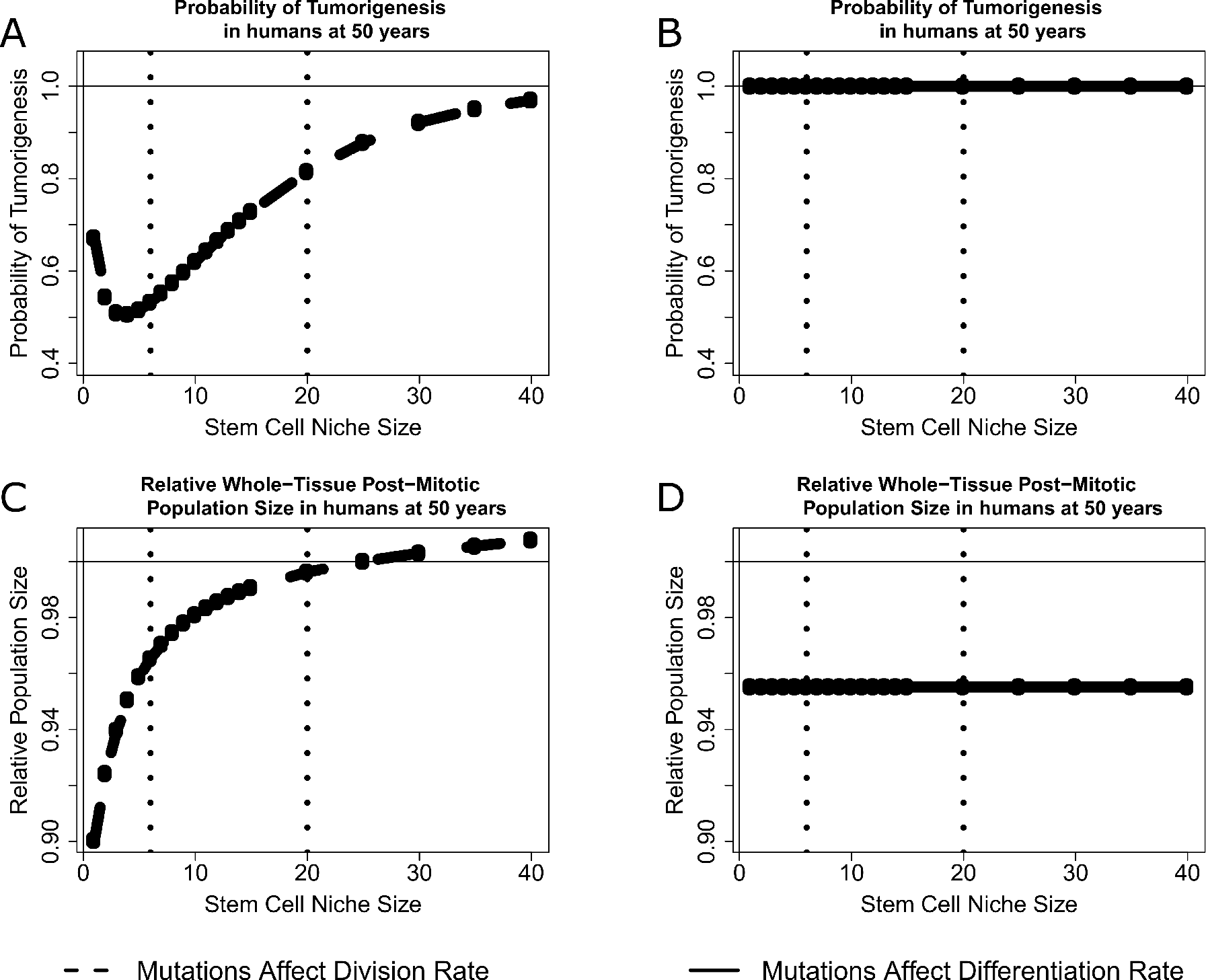
An Evolutionary Tradeoff with Niche Size in Humans. For scenarios where mutations affect division rate, tumorigenesis is minimized at an intermediate population size (A), at the expense of fixing deleterious mutations and decreasing tissue renewal (C). For scenarios where mutations just affect differentiation rate, tumorigenesis is minimized for large population sizes (B), and the effect of mutation accumulation on the epithelium is independent of niche size (D). The black vertical dotted lines are placed at 6 and 20, the reported stem cell population sizes in humans.

## 4 Discussion

**The effects of mutational target on optimal stem cell niche size**. When mutations confer a selective advantage or disadvantage within the niche, there exists an intermediate crypt size that minimizes the probability that any crypt accumulates the large beneficial mutations necessary to initiate a tumor. This result reinforces previous results by Michor et al. [2003], who also found an intermediate niche size exists that decreases the rate of tumor initiation for mutations under selection. At small stem cell niche sizes there exists a large number of crypts to maintain homeostasis, and a higher probability that any one crypt will obtain a rare mutation of large effect that would result in tumorigenesis. As stem cell niche size increases, the number of crypts needed to maintain the same amount of epithelium decreases, and so does the probability of fixing mutations within the crypts, and therefore the chance of fixing a rare mutation of large effect. However, for larger values of stem cell niche size, the strength of selection increases, thus increasing the chance that a fixed mutation was beneficial, leading to higher chances of tumorigenesis. Here, given the specified parameters for mice, we find that the minimum probability of tumorigenesis exists at 7 cells, which is within the empirically reported stem cell number [Kozar et al., 2013, Ritsma et al., 2014] for intestinal stem cell niches in mice. At this population size the whole tissue size is expected to decline with age as deleterious mutations accumulate in stem cell niches, and if selective pressures against tumorigenesis have selected for stem cell niche population sizes at this intermediate then it has been at the expense of increasing epithelial attrition. When considering parameters reported for the human intestinal epithelium and mutations that affect division rate, we find that the population size with the smallest probability of tumorigenesis is lower than the reported sizes. We note that the experimental methods used to elucidate stem cell population size in mice are not possible in humans, and understanding the true size of the stem cell niche in larger, long lived animals will shed light on how long lived organisms balance maintaining their tissues while minimizing mutation-associated disease.

When mutations affect differentiation rate, and thus fix neutrally in the stem cell niche, larger stem cell niche sizes result in a lower probability of tumorigenesis. However, the rate at which the epithelium changes total size as mutations accumulate within stem cell niches is independent of stem cell niche size. At small niche population sizes, the number of fixed mutations in the entire system is maximized as both the number of crypts and probability of fixation of every mutation within each crypt is maximized, thus leading to the maximum probability that any one fixed mutation resulted in tumorigenesis. Because these mutations fix neutrally, the expected value of a fixed mutation is independent of stem cell niche population size. Furthermore, the probability of fixation of a mutation in a niche is the inverse of the contribution of that niche to tissue homeostasis. For instance, a mutation that arises within a niche that is ten times the size of another niche has ten times smaller probability of fixation, but ten times higher influence on the total epithelium, meaning that the expected influence of the total amount of accumulated mutations in systems with different stem cell population sizes but consistent total epithelium sizes is invariant.

It is probable that the mutational target size for mutations that affect the propensity for stem cells to commit to differentiation is smaller than that for mutations that might affect the overall division rate, and therefore selection may not act as strongly to optimize niche size in light on minimizing tumorigenesis caused by failure to differentiate. Furthermore, our model results indicate that both the probability of tumorigenesis and the extent of tissue size change are larger for the scenarios where mutations only affect differentiation rate. This is due to mutations in this scenario fixing neutrally and having a larger absolute influence towards tumorigenesis (we model expected mutational effect sizes as a proportion of the rate they govern). We use the same rate of mutation when modeling both mutations that affect division rate and differentiation rate, but given that there is probably a smaller target size for mutations that change the rate at which stem cells commit to differentiation, there will also be a smaller mutation rate.

**The intestinal epithelium population is expected to decline with age through stem cell attrition**. When employing distributions of mutational effects commonly found in experiments on whole organisms we find that the total intestinal epithelium size is expected to decrease with age. This attrition is modest in the mouse, with a mouse three years into adulthood having an intestinal epithelium approximately 0.3% – 0.5% smaller than a mouse that just entered adulthood. This attrition is potentially much more substantial for humans, given their longer lifetime. When mutations affect division rate and for the scenario where the stem cell niche size exists as 20 cells, we find remarkably similar total epithelial population change to the mouse scenario because, despite the longer lifetime, the division rate of stem cells is slower and the population size is larger, decreasing the strength of drift and the accumulation of deleterious mutations. When mutations fix neutrally for the same population size, and affect differentiation rate, we calculate that the total post-mitotic epithelium will decrease approximately 5% at 75 years post-adulthood. If the true stem cell niche size in humans is closer to the lower estimate of 6 cells, we calculate a much higher rate of tissue attrition, with the epithelium decreasing over 10 and over 20 percent in the division and differentiation scenarios, respectively. This is due to the much stronger influence of deleterious mutations drifting to fixation at smaller population sizes.

**Mechanisms of intestinal homeostasis and evolutionary tradeoffs**. Although mice and humans have an intrinsic rate of crypt division, this event is exceedingly rare [Snippert et al., 2014, Baker et al., 2014] and likely does not play a role in maintaining tissue homeostasis. Thus, population equilibrium during normal tissue maintenance is primarily controlled through the continual division and differentiation of stem cells. Here, we show that, due to the small population sizes of the independently evolving populations of stem cells constituting the intestinal epithelium, the total tissue size is expected to decrease over an organism’s lifetime due to the accumulation of deleterious mutations in stem cell compartments. Furthermore, we have demonstrated that the intestinal stem cell compartment is an example of Muller’s ratchet and mutational meltdown, or, the gradual decline in fitness and population size associated with small asexual populations [Lynch et al., 1993]. These small compartments minimize the overall probability that any mutation in the intestines results in the origin of a tumor when mutations confer a selective advantage in the niche, and is thus a prime example of an evolutionary tradeoff.

